# Uncompetitive Allosteric Inhibitor of Mitochondrial Creatine Kinase Prevents Binding and Release of Creatine by Stabilization of Loop Closure

**DOI:** 10.64898/2026.01.08.696683

**Authors:** Merve Demir, Chen-Ting Ma, Nikhil Puvvula, Laura Koepping, Palak Gosalia, Sirkku Pollari, Shaun Alba, Zoe Zong, Ya Li, Lynn Fujimoto, Andrey Bobkov, Taro Hitosugi, Jianhua Zhao, Eduard Sergienko

## Abstract

Mitochondrial creatine kinase (MtCK) is a key enzyme in energy buffering and homeostasis in cells. It catalyzes transfer of phosphoryl group from ATP to creatine. Overexpression of MtCK occurs in many cancer cells to meet elevated energy demands, which is associated with poor prognosis. This suggests that MtCK may be a promising target for cancer therapeutics. We sought to discover first-in-class selective inhibitors of MtCK with diverse mechanisms of action using high-throughput screening, biochemical characterization and cryo-EM studies. Through these studies, we identified diverse types of compounds that modulate activity of MtCK *in vitro*, including fast-equilibrium and time-dependent orthosteric and allosteric inhibitors. Select hits were subjected to *in vitro* enzymatic and binding assays to assess MtCK inhibition and binding. A subset of inhibitors was advanced into structural studies using cryo-EM resulting in the molecular structure of MtCK with and without bound substrates in complex with an allosteric uncompetitive inhibitor that we discovered. These studies identified compound’s unique binding pocket on MtCK and established the molecular steps manifesting in the apparent uncompetitive mode of inhibition. We demonstrate that the compound stabilizes an active site loop in a closed conformation restricting access of the creatine substrate to and the release of phosphocreatine product from its binding pocket. These findings establish chemical tools suitable for validation of MtCK as a promising breast cancer target with high therapeutic potential and build a foundation for future structure-guided optimization of the hit compounds of MtCK we identified and de novo rational design of novel MtCK inhibitors.

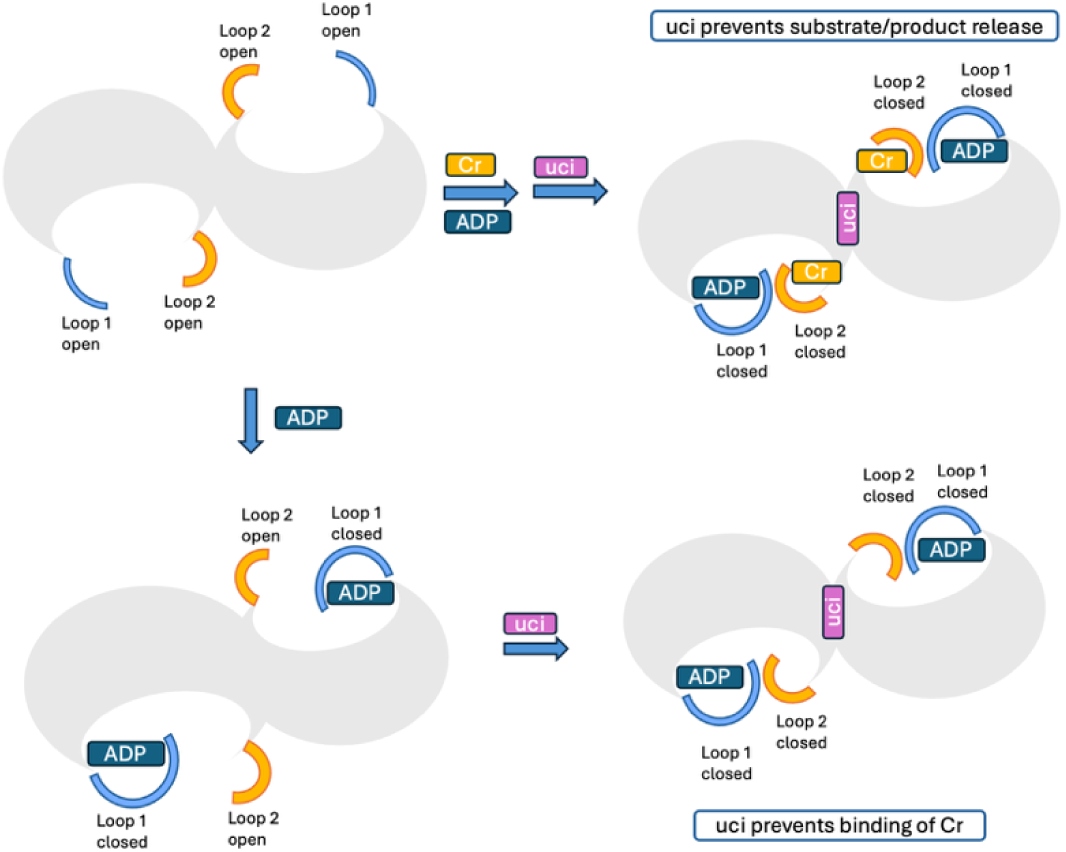

## Introduction

Mitochondrial creatine kinase (MtCK), an enzyme involved in energy supply and homeostasis in cells, has emerged as a potential cancer target implicated in tumor metabolism and proliferation[1–7]. MtCK1 located in the mitochondrial intermembrane space, functions in a key step in the phosphocreatine (pCr) energy shuttle. It transfers a phosphoryl group from mitochondrial ATP to creatine, which produces phosphocreatine that can be converted back to creatine by cytosolic creatine kinase (CK) to regenerate ATP for buffering energy demands. Recent studies have highlighted the addiction of cancer cells to MtCK enzyme that provides enhanced ATP buffering and supply under high energy demands [2,8,9]. Elevated expression of MtCK has been observed in various cancer types including HER2+ breast cancer and EVI+ AML cells, suggesting its potential role in tumor progression [8–10]. According to the previous study [9], HER2 signaling enhances the pathway by inducing ABL-dependent phosphorylation of MtCK1 at Y153, which stabilizes MtCK1 and increases pCr energy shuttle flux. As a result, phosphocreatine production by MtCK1 increases ATP regeneration in the cytosol to support rapid tumor proliferation in HER2^+^ breast cancer (**Figure 1**).

**Figure 1.**
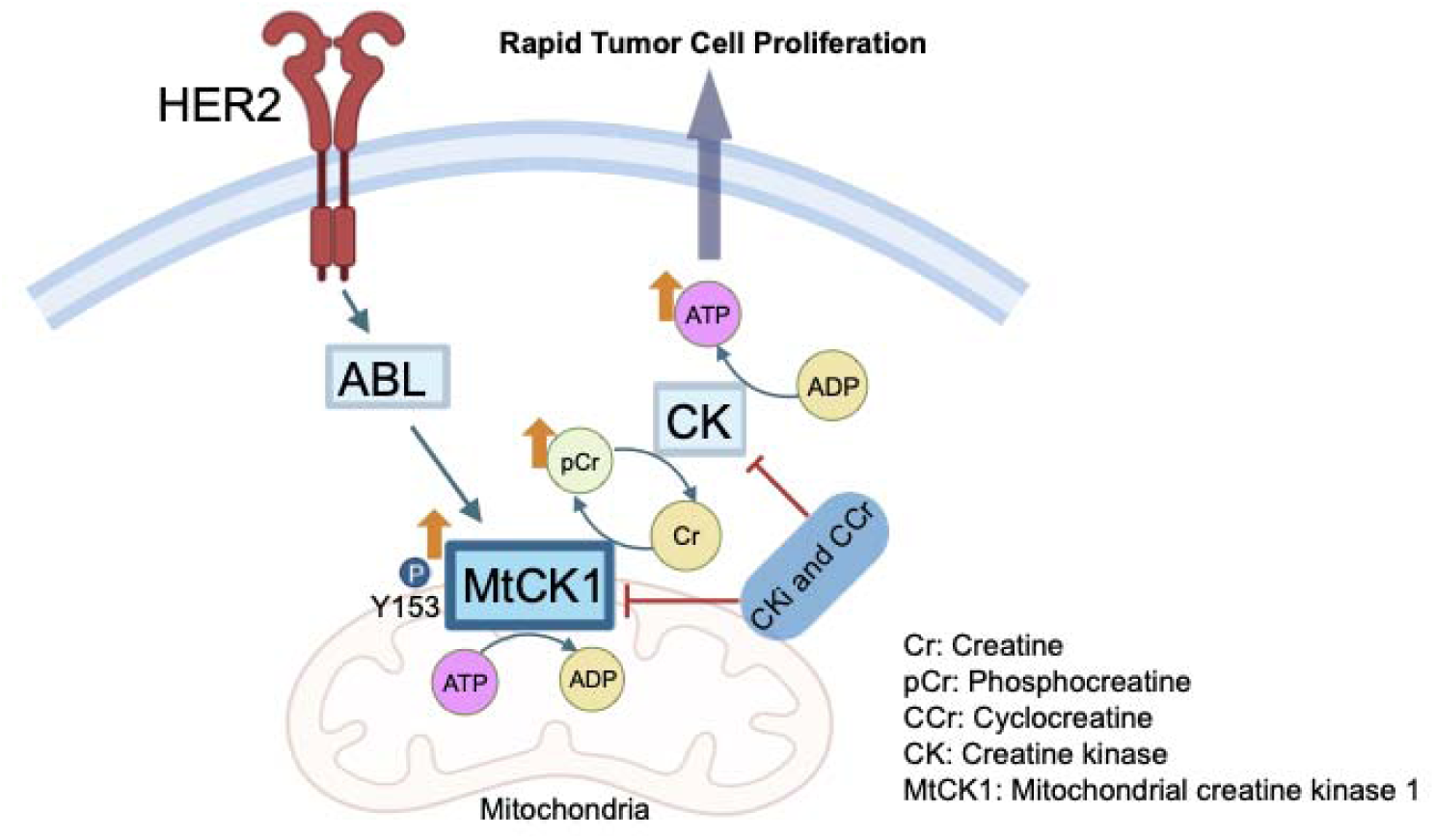
Depiction of role of MtCK1 in HER2 cancer biology

Breast cancer is one of the leading causes of cancer-related mortality among women worldwide despite advances in early detection and targeted therapies. Patients of HER2^+^ breast cancer frequently develop resistance to HER2 targeted therapies including trastuzumab and pertuzumab [11]. Therefore, targeting downstream pathway of HER2 signaling such as MtCK1 offers a new strategy to overcome HER2 targeted resistance and improve the outcomes of HER2^+^ breast cancer patients. Inhibition of MtCK can disrupt metabolic adaptation of cancer cells, rendering them more susceptible to apoptosis and cell death. Despite apparent importance in tumor metabolism, MtCK has not yet been thoroughly investigated and no selective inhibitors were reported to date. The lack of specific chemical probes have limited the functional interrogation of MtCK in cancer models and hindered the development of MtCK-targeted therapeutics. Only a few MtCK inhibitors targeting its active site exist, such as cyclocreatine (cCr) and creatine kinase inhibitor (CKi), however, both are nonselective and inhibit both mitochondrial and cytoplasmic creatine kinase enzymes [9,11]. At the same time, these compounds clearly establish the importance of the phosphagen system driven by MtCK for cancer cell survival [9,11,12]. Addressing this gap, we employed a comprehensive strategy that integrates high-throughput screening (HTS), biochemical and structural validation, and computational modeling to identify novel small-molecule inhibitors with therapeutic potential.

In this study, we screened a broad collection of small-molecule libraries to identify compounds that inhibit enzyme catalytic activity. Selected hit compounds were further evaluated *in vitro* enzymatic and binding assays using MtCK wild type and mutants incorporating modifications of specific active site residues to assess their inhibition and binding mode [12]. Unsurprisingly, most competitive inhibitors expected to occupy the MtCK active site were affected by these mutations. Interestingly, the uncompetitive inhibitor of the wild-type MtCK identified through our screening efforts stimulated the catalytic activity of the MtCK mutants that had creatine binding defect. We obtained structures of MtCK in complex with the uncompetitive inhibitor with and without substrates. Comparison of these structures suggested that the uncompetitive inhibitor stabilizes the flexible loop that forms the creatine binding site and exert its effects on binding of creatine (Cr) and overall MtCK catalytic reaction. These findings help to explain the inhibitor’s mechanism of action and lay the groundwork for optimization of the inhibitor scaffold and advancing to cellular validation and preclinical development and may open new avenues for therapeutics targeting mitochondrial function in cancer.

## Methods and Materials

### Protein Production

The pet15b vector containing human uMtCK gene and single point mutations of uMtCK were created as previously described [13]. Rosetta 2 (DE3) cell lines were transformed with pet15b vector containing human uMtCK or its mutants and proteins were overexpressed with 0.5 mM IPTG overnight at 15 C. Cell pellet resuspended in lysis buffer (50 mM Tris-HCl pH 8.0, 200 mM NaCl and 5% glycerol) was supplemented with protease inhibitor cocktail, DNase1 and 10 mM MgCl_2_ and lysed by sonication. After a spin down, supernatant containing protein was purified using HisPur Co-NTA beads using elution buffer containing 50 mM Tris-HCl pH 8.0, 250 mM NaCl, 500 mM imidazole and 10% glycerol, which is followed by size exclusion column HiLoad 26/600 Superdex 200. The peak for octameric protein were collected in a final buffer condition of 50 mM Tris-HCl pH 8.0 and 200 mM NaCl, and the aliquots were stored at –80 C.

### Activity Assays for Screening

An assay buffer containing Tris pH 8.5, 7 mM MgCl_2_, 250 mM sucrose, 2 mM DTT and 0.005% Tween 20 was used to prepare enzyme (MtCK) and substrate (ATP and creatine) dilutions for high-throughput screening and dose-response experiments in 1536-well plate. For 1xK_m_ conditions, final assay concentrations of 0.25 mM ATP and 0.8 mM creatine (Cr) were utilized. Optimal MtCK enzyme assay concentration was 0.3 nM. For 20xK_m_ conditions, the final assay concentrations of ATP and Cr were 5 mM and 16 mM, respectively. Enzyme was dispensed to compound containing plates with 0 or 2 hours incubations. Substate stock solution was dispensed to plates and the reaction mixtures were further incubated for 30 min. The amount of the generated concentration of ADP product was measured using Promega ADP-Glo Kit following manufacturer protocol. Plates were read on PHERAstar FS microplate reader to measure the luminescence output. The PGK assay was utilized as the counter screen for detection of compounds that interfere with the detection system and thus result in false positives. Similarly, an assay buffer of HEPES pH 7.2, 10 mM MgCl_2_, 0.1 mg/ml, 1 mM DTT and 0.005% Tween 20 was prepared for dilutions of phosphoglycerate kinase (PGK) and substrates, ATP and 3-phosphoglycerate (3PG). GAPDH and NADH were included in the PGK reaction to promote the direction of ADP generation. For 1x and 20xK_m_ ATP conditions, a final concentration of 0.07 and 1.4 mM ATP were utilized in the presence of 10 mM 3PG, 1.5 uM GADPH and 0.5 mM NADH and final PGK concentration of 0.5 nM. Data were analyzed using GraphPad Prism.

### Microscale Thermophoresis

The WT and mutant uMtCK were buffer exchanged to 50mM phosphate, pH 8.0 containing 150 mM NaCl and incubated with Dylight 650 NHS ester for 1 hour at room temperature on rocker at dark for conjugation reaction as previously described [13]. The reaction products were dialyzed to the storage buffer (50 mM Tris-HCl pH 8.0 and 150 mM NaCl) and the aliquots of labeled proteins were stored at –80°C in the dark.

For the binding assay, compounds were serially diluted using assay buffer containing 50 mM Tris-HCl pH 8.5, 5 mM MgCl_2_, 100 mM NaCl, 5% DMSO and 0.05% Tween 20 and mixed with equal volume of protein. The samples were run on NanoTemper Monolith NT.115 MicroScale Thermophoresis instrument at 22 °C to measure the amount of unbound and bound protein across varying concentration of compounds and data was fit into a single binding isotherm to extract the K_d_ value.

### Cryo-EM Data Collection and Processing

Cryo-EM sample preparations and data collection were performed in the Sanford Burnham Prebys Cryo-EM Core facility as previously described [13]. 4ul of 0.5-1 mg/mL of uMtCK protein was applied to a Quantifoil holey carbon grid (R2/2, 300mesh, copper) that was glow-discharged with a Pelco easiGlow at 15 mA for 25 s. The sample was blotted using Vitrobot MarkIV for 10 s at 4 °C, 100% humidity and blot force 0 and plunge-frozen in liquid ethane. For samples with substrates and uncompetitive inhibitor, uci, protein was preincubated with 10 mM Cr and/or 1 mM ADP, 50 mM NO_3_, 50 mM MgCl_2_ and 0.5 mM uci for 15 minutes.

Grid screening and data collection was performed in a Titan Krios microscope (Thermo Fischer) at 300kV, using SerialEM [14] by image shift (four shots per hole) with beam-tilt compensation. Movies with 20 frames each were collected using a Gatan K3 detector in super resolution mode with a dose rate of around 20 electrons per pixel per second and with a total electron dose of around 34 e-/A2, at a pixel size of 1.06A per physical pixel. All micrographs were collected with a defocus ranging between 0.9 to 2.1 μm. Movies were processed in Cryosparc 3.3.2 [15] by motion-correction and dose-weighting of frames by Patch motion correction using default parameters (**Figure S1**). The CTF parameters were estimated by Patch CTF estimation and binned by 2. Images with ice contamination, failed CTF estimation or low resolution CTF fit values were discarded with Manually Curate Exposures. Particle picking was done with a template picker generated using previous MtCK density maps with a round blob with particle diameter of 145 A and extracted with a box size of 256 pixels and binned by 4. The binned particles were cleaned by 2D classification. Unbinned particles were used to generate an Ab-Initio reconstruction with D4 symmetry enforced, followed by a 3D reconstruction using Non-uniform Refinement.

The cryo-EM structures, 9B05 and 9B14, were used as a starting model and was refined using an iterative process of manual adjustments and refinement using Coot v.0.9.8.96 and Phenix v.1.21.2 [16,17]. The ligand restraints files for ADP, CRN and A1CZQ for uci were generated using eLBOW in Phenix with 3-letter codes of the ligands available in PDB or SMILES string for uci and eLBOW AM1 QM optimization method. The final restraints and metal ion coordinations were generated in ReadySet by providing the CIF files generated in eLBOW. Finally, the model PDB file and generated restraints CIF files were used in the refinement of the MtCK structures with phenix.real_space_refine. The final model was validated using MolProbity and Phenix. The resulting cryo-EM maps and models were visualized using UCSF ChimeraX [18].

## Results and Discussion

### MtCK Inhibitor Identification

Previously identified inhibitor of CKs (CKi) showed limited efficacy against breast cancer cells in cellular assays despite the promising *in vitro* data and ability to covalently block the active site of both mitochondrial and cytoplasmic creatine kinases [12,13]. We set out to identify small molecule modulators of MtCK with a diverse mechanism of action using high-throughput screening of broad selection of commercially available small molecule libraries. The activity assay in forward direction using Promega ADP-Glo kit was optimized for screening in 384- or 1536-well plate formats. Compounds were tested at a single concentration of 25 µM at 1xK_m_ concentrations of creatine (Cr) and ATP. Small molecules showing inhibition were retested at 25 µM using triplicate wells. The PGK/ADP-Glo assay was employed to eliminate false positives that inhibited the detection system. Identified MtCK-specific hits were purchased as dry powders and confirmed in dose response curves using 16-compound concentration duplicates. 1x and 20xK_m_ conditions for both ATP and Cr substrates were used to assign compounds into competitive, noncompetitive and uncompetitive MtCK inhibitor apparent groups. IC_50_ values from 20xK_m_ condition much higher than IC_50_ from 1xK_m_ condition indicated competitive inhibitors, while the reverse indicated uncompetitive inhibitors and similar IC_50_ values provided the noncompetitive inhibitors group. In addition to no pre-incubation assays, these assays were performed with 2-hour pre-incubation of compounds with the protein to distinguish rapid-equilibrium and time-dependent inhibitors. As a result of these extensive profiling studies, we identified two competitive (**cmi-1 and -2**), two time-dependent (**tdi-1 and -2**), one uncompetitive inhibitor (**uci**) and one activator (**act**) scaffold of MtCK with IC_50_/EC_50_ values lower than 15 µM shown in **Figure 2A** and **Table 1**. Although, both uci and act compounds demonstrate IC_50_/EC_50_ values ∼100 µM in 1xK_m_ assay, their potency at 20x K_m_ condition were 4 and 14 µM, respectively.

**Figure 2.**
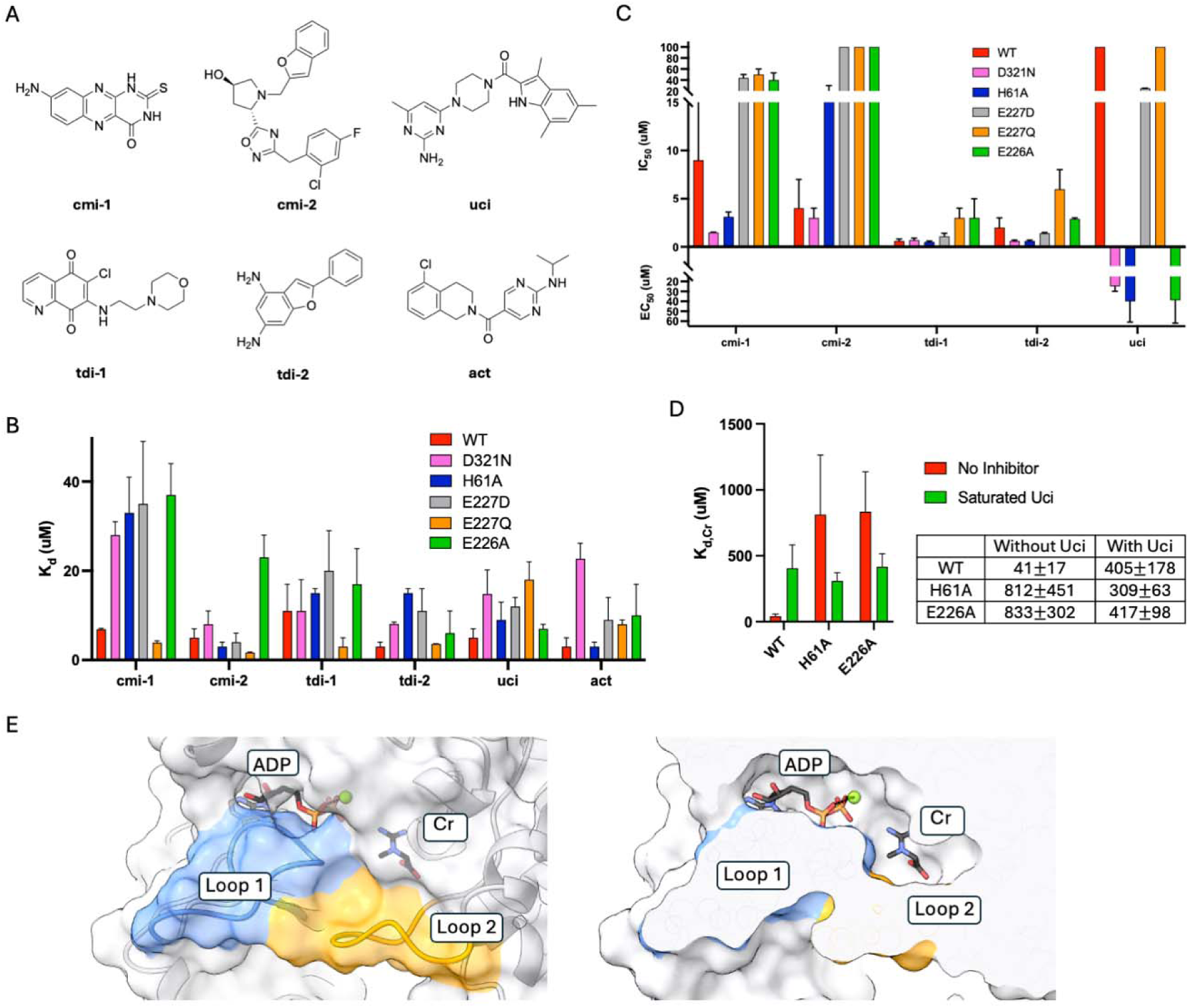
A) The structures of inhibitors of MtCK with varying mechanism of action. B) Bar graph of dissociation constants, K_d_, of each inhibitor with WT and mutant MtCK. C) Bar graph of inhibitors with WT and mutant MtCK represent the results of dose response experiments with IC_50_ and EC_50_ values with 2 hour incubation. D) Bar graph and table of binding data for Cr with WT, H61A and E226A with or without uci. Data shown as mean ± SD from at least three independent experiments. E) Active site view of MtCK in complex with transition state analogs, ADP and Cr, (PDB ID: 9B14) shows flexible loops that interlock to form the substrate binding pocket, shown as both surface view and cross-section of the catalytic pocket.

**Table 1.** IC_50_ or EC_50_ values of inhibitors with WT or mutant MtCK^a^.

| IC<sub>50</sub> or EC<sub>50</sub> (μM) |  | cmi-1 | cmi-2 | tdi-1 | tdi-2 | uci | act |
| --- | --- | --- | --- | --- | --- | --- | --- |
| 1xK<sub>m</sub> | WT | 5±3 | 4±3 | 0.6±0.2 | 1.4±1.3 | >100 | >100 |
|  | D321N | 1.48±0.04 | 3±1 | 0.7±0.2 | 0.6±0.1 | 25±5<sup>b</sup> | >100 |
|  | H61A | 3.1±0.5 | 18±12 | 0.5±0.1 | 0.6±0.1 | 40±21<sup>b</sup> | >100 |
|  | E227D | 38±10 | >100 | 1.1±0.3 | 1.4±0.1 | 23±4 | >100 |
|  | E227Q | 44±12 | >100 | 4±1 | 5±2 | >100 | >100 |
|  | E226A | 40±13 | >100 | 3±2 | 2.9±0.1 | 39±23<sup>b</sup> | >100 |
| 20xK<sub>m</sub> | WT | 17±4 | 67±28 | 0.39±0.04 | 0.5±0.1 | 3.9±2.5 | 14<sup>b</sup> |
|  | D321N | 3±1 | 56±25 | 0.4±0.1 | 0.50±0.02 | >100 | 65±13<sup>b</sup> |

<sup>a</sup> Values are reported as mean ± SD for dose response experiments with 2 hour incubation from at least three independent experiments. <sup>b</sup> Activation potency EC<sub>50</sub> is reported for these values, while the rest of the reported values are inhibition potency IC<sub>50</sub>.

### Compound Binding and Inhibition with MtCK Mutants

To confirm the binding of small molecule modulators by MtCK, we performed binding assays using microscale thermophoresis and determined the binding constant K_d_ for each compound with WT MtCK. The binding affinities of the compounds were in the range of 3-12 µM (**Figure 2B and Table 2**). Overall, the compounds demonstrated binding affinity that matches their inhibition or activation potency. We next evaluated the binding affinities and inhibitory potencies of the compounds using previously designed and characterized MtCK mutants, D321N, H61A, E227D, E227Q and E226A, aiming to assign the specific binding locations for the small molecules. As previously described [12], we strategically selected these residues since Glu227 and Glu226 showed their importance for catalytic activity, while loop residues Asp321 and His61 appeared to have a role in substrate binding[19–23]. MtCK possesses two substrate-binding pockets and two corresponding flexible loops that shield the pockets and interlock upon binding of both substrates providing optimal environment for catalysis (**Figure 1E**). The mutations within each of the two loops or binding pockets are expected to alter the binding affinity of the small molecules, thereby providing insight into their potential binding locations.

**Table 2.** Binding data results for inhibitors with WT and mutant MtCK^a^.

| K<sub>d</sub> (μM) | WT | D321N | H61A | E227D | E227Q | E226A |
| --- | --- | --- | --- | --- | --- | --- |
| cmi-1 | 6.9±0.2 | 28±3 | 33±8 | 35±14 | 3.9±0.4 | 37±7 |
| cmi-2 | 5±2 | 8±3 | 3±1 | 4±2 | 1.7±0.1 | 23±5 |
| tdi-1 | 11±6 | 11±7 | 15±1 | 20±9 | 3±2 | 17±8 |
| tdi-2 | 3±1 | 8.1±0.4 | 15±1 | 11±5 | 3.6±0.1 | 6±5 |
| uci | 5±2 | 14.8±5.4 | 9±4 | 12±2 | 18±4 | 7±1 |
| act | 3±2 | 22.7±3.5 | 3±1 | 9±5 | 8±1 | 10±7 |

<sup>a</sup> Dissociation constants, $K_d$ , are reported as mean ± SD from at least two independent trials.

Competitive inhibitor 1 (cmi-1) showed a 4-5 fold decrease in binding affinity with mutations at D321, H61, E227 and E226, however only mutations of E227 and E226 residues affect inhibitory potency in the MtCK activity assay as shown in **Figure 2B&C and Table 1&2,** indicating that cmi-1 is likely binding in the active site. Cmi-2 had showed no change of inhibition for D321N and only a moderate loss of inhibitory potency with mutation H61A, while all mutations of E226 and E227 abolished inhibition, suggesting that this inhibitor may bind in the vicinity of and directly interact with these residues.

Time-dependent inhibitor 1 (tdi-1) exhibited similar affinity and inhibition levels with all mutants showing only a minor reduction of potency for E227 residue mutations. A conservative E227D substitution resulted only in 2-fold decrease, while substitutions with loss of the negative charge (E227Q) showed 7-fold decrease of inhibition potency; similar effect had neutralizing substitution at E226 residues. Interestingly, the fact that the IC_50_ values at 1xK_m_ and 20xK_m_ conditions were similar suggests that the inhibitor binding is insensitive to substrate presence, i.e noncompetitive, and thus may take place at an allosteric site, explaining the lack of effect from active site mutations. On the other hand, tdi-2 displayed a moderate decrease in binding with H61A and E227D, while its inhibitory potency lowered by 3-fold with E227Q that implies that tdi-2 may be binding by Cr binding site.

Uncompetitive inhibitor (uci) and activator (act) of MtCK are expected to occupy allosteric sites; consistently they showed similar binding affinities with different active site mutants. Interestingly, act displays a significant decrease in binding by replacement of D321 with asparagine. Similarly, the activity assay at 20xK_m_ condition with D321N showed decreased levels of activation with act. These results suggest that act may be binding by the ATP loop or binding pocket near D321 and aiding efficiency of binding of substrate ATP that results in increase of enzyme activity. We were unable to test other mutants at 20xK_m_ condition due to the need of high concentrations of protein or substrates that are not practically attainable for these mutants that either have high K_m_ values for Cr or slow reaction rates. However, our activity assay at 1xK_m_ revealed an interesting result with uci where we observed activation of reaction with the mutants D321N, H61A and E226A that we have shown to demonstrate Cr binding deficiency [12]. Uncompetitive inhibitors bind in the presence of substrates and slow down the overall reaction or prevent the release of products. With MtCK mutants with Cr binding defect, uci appears to assist Cr binding that generates increased levels of product. We next measured the binding affinity for Cr in the absence and presence of uci with WT, H61A and E226A as shown in **Figure 2D**. In the presence of uci, WT MtCK significantly loses affinity for Cr, while H61A and E226A showed a moderate increase in affinity for Cr that supports our findings from the activity assays. To get a deeper understanding of these findings and establish the binding site of uci with respect to Cr binding, we obtained cryo-EM structure of WT MtCK bound to the compound with or without substrates.

### Structural Modeling of uci in complex with MtCK

Uncompetitive inhibitors bind their target protein allosterically and prevent the catalytic conversion of the substrates and release of products. To understand the mechanism of binding and inhibition of uci at a molecular level, we determined the structures of WT MtCK incubated with uci (uci-MtCK) alone, with both TSA and uci (uci-TSA-MtCK) or with ADP and uci (uci-ADP-MtCK) present. The data collection and refinement statistics are presented in **Table S1**. MtCK is an octamer composed of four equivalent homodimer units. All structures showed additional density between two monomers of each homodimer of MtCK that could fit uci as a ligand (**Figure 3A&B**). For simplicity, we will show the fit of different orientations of uci in the density map of uci-TSA-MtCK complex.

**Figure 3.**
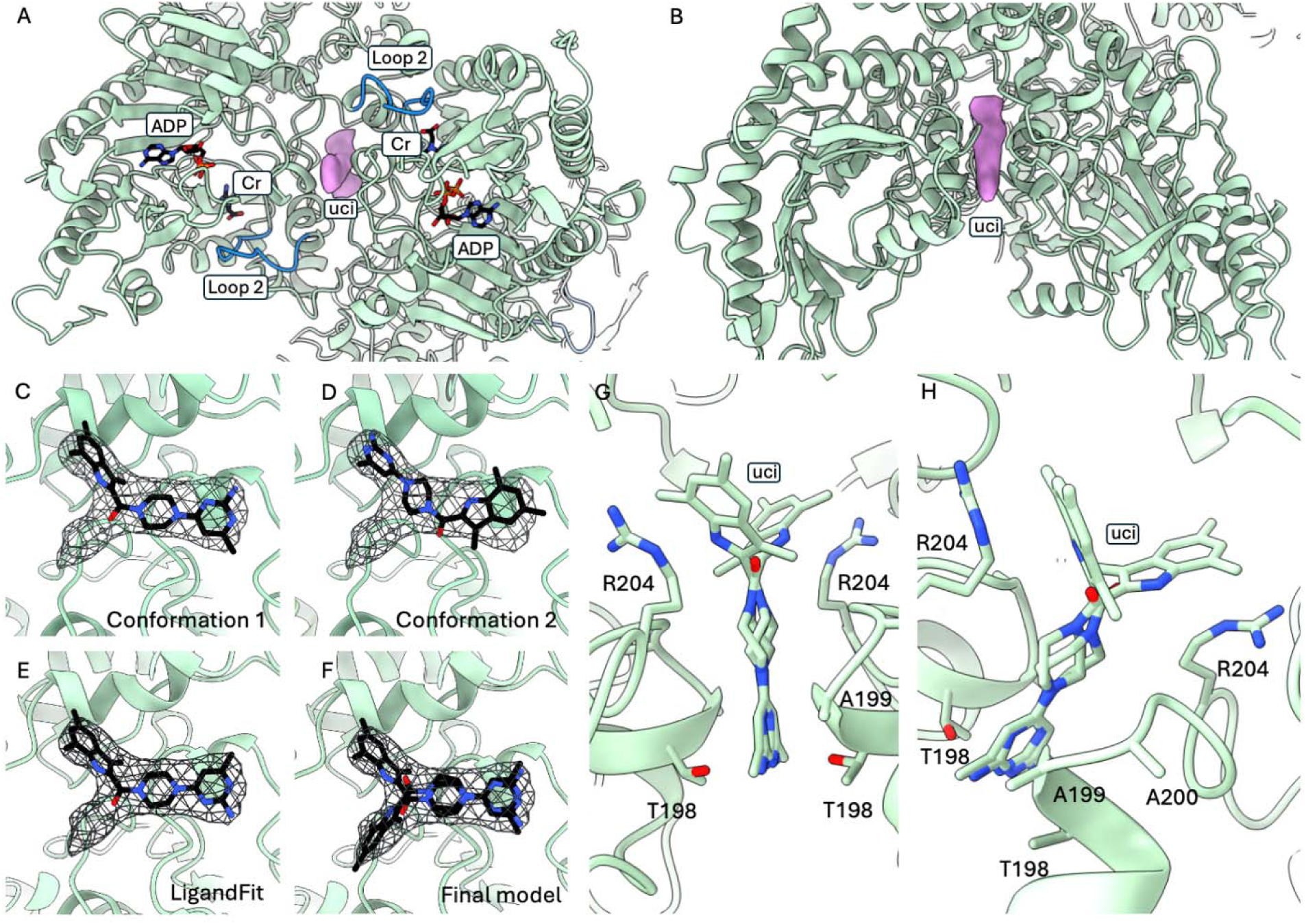
A) Difference map, calculated using the density map obtained for uci-TSA-MtCK and previously obtained TSA-MtCK structure (PDB ID 9B14), shows the additional density shown in pink where the uncompetitive inhibitor, uci, binds in close proximity to Cr binding site and Loop 2. B) Side view of the homodimer with the density for uci is shown. C) Manual fit of conformation 1, D) manual fit of conformation 2, E) resulting fit from LigandFit from Phenix tools and F) final model of the two copies of uci into the density are shown. G) Pyrimidine group of uci is stacked between residues 198-200 from each monomer of the homodimer. H) Residue R204 forms a cation-Jr interaction with indole group of uci.

We have modeled two conformations of uci with a 180° flip that fit in the observed density to identify the best fit of the ligand **Figure 3C&D**. Due to the symmetry of the protein, we observe density for two copies of the ligand corresponding to two equivalent poses of the ligand. Initial models are shown with only one copy modeled to evaluate the fits more clearly. Conformation 1 places the indole group near N-terminal domain and pyrimidine group near the Cr binding site and Loop 2 that closes over the Cr binding site. Conformation 2 of uci places the indole group near the Cr binding site and Loop 2. After a round of refinement and visual inspection, conformation 1 appears to fit better into the density than the conformation 2. Next, we used LigandFit tool in Phenix to find the best fitting orientation of uci into the density observed for the compound. LigandFit was set up with density map, ligand free model, PDB file of ligand uci, number of copies of ligand set to 4 and number of conformers set to 10. This resulted with conformation 2 favored in all four sites observed for uci ligand (**Figure 3E**). We used this conformation to model the rest of the structures. Finally, as the compound can bind to either monomer of each homodimer, we observe density for two copies of uci that were modeled as alternate conformations of each other to avoid clashes as shown in **Figure 3F**. In this conformation, pyrimidine group of uci is stacked between residues T198-A200 of each monomer of the homodimer (**Figure 3G**) while indole group forms a cation-Jr interaction with R204 (**Figure 3H**).

### Binding pocket and mode of uci

The comparison of uci-MtCK structure to unliganded-MtCK showed no conformational changes in the protein upon binding to uci (**Figure 4A**). Similarly, we have not observed any additional conformational changes in uci-TSA-MtCK in comparison to TSA-MtCK structure (**Figure 4B**). However, uci binds in a pocket located near the Cr binding site. Consistent with this, activity and binding assays had suggested that uci may influence Cr binding and release. In the uci-TSA-MtCK structure, the presence of both substrates prevents observation of the inhibition effect of uci, as the substrates remain in their bound conformations if uci is present as shown in **Figure 4**. Conversely, removal of both substrates offers limited insight, since the relevant loops are flexible and only stabilize upon substrate binding.

**Figure 4.**
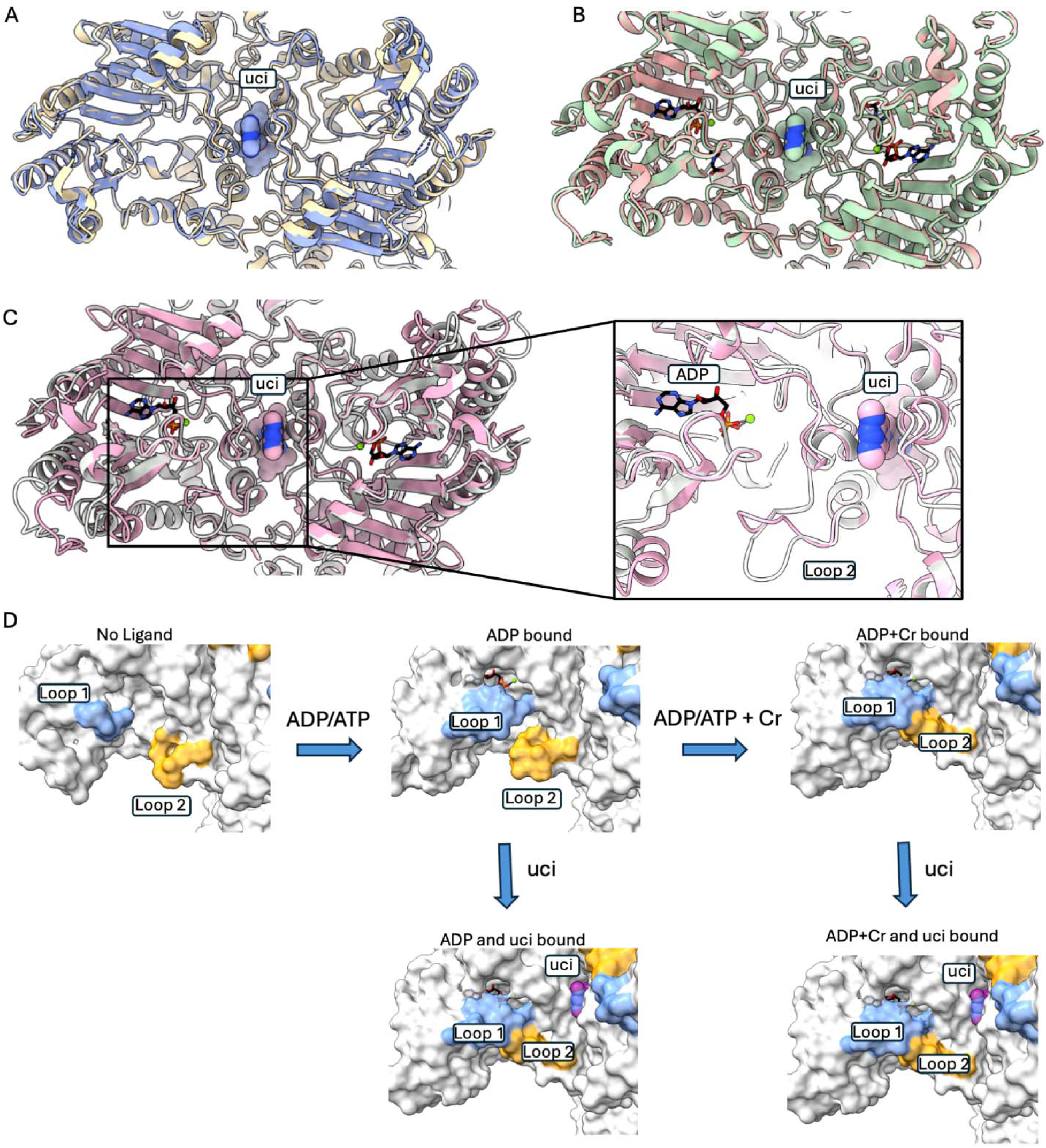
A) Overlay of uci-MtCK (in corn blue) with unliganded-MtCK (PDB ID 9B05, in pale yellow) did not indicate any conformational change upon uci binding B) uci-TSA-MtCK (in green) aligned with TSA-MtCK structure (PDB ID 9B14, in pale pink) had no changes with uci present C) Alignment of uci-ADP-MtCK (in pink) with ADP-MtCK (PDB ID 9B04, in white) showed that Loop 2 adopts a closed conformation upon uci binding in the absence of Cr. D) A structural representation of the proposed mechanism of inhibition of the uncompetitive inhibitor of MtCK. In the unliganded MtCK, loops 1 and 2 are disordered or in open conformation. Upon ADP binding, Loop 1 closes over the active site while Loop 2 remains open in the absence of Cr, however uci binding promotes Loop 2 closure. When both substrates are bound, both loops are closed over active site and remains interlocked in the presence of uci.

Since closure of Loop 2 depends on the closure of Loop 1 and Cr binding [13], we next obtained the structure of MtCK bound to uci and ADP (uci-ADP-MtCK) to assess whether uci promotes closure of Loop 2 in the presence or absence of Cr that can shed light into the inhibition mechanism of uci. Strikingly, in the absence of Cr, Loop 2, which remains predominantly in open conformation in ADP-MtCK [13], adopts its closed conformation when uci is bound (**Figure 4C**). Interestingly, residues L196 and L197 also adopt similar conformations to what is observed only in the presence of Cr in uci-ADP-MtCK structure. The loop closure in the presence of uci is consistent with the loss of affinity for Cr in the binding assays with WT MtCK as it would impede Cr access to the active site. Furthermore, once the substrates are bound, loop closure appears to lock substrates in place and prevent the product release. In summary, uci stabilizes the closed conformation of Loop 2 in the presence of ATP/ADP, thereby interfering with Cr binding and phosphocreatine product release and hereby inhibiting the enzymatic reaction (**Figure 4D**). On the other hand, alanine substitution of H61, a residue that is located in Loop 2 and required for its interlock with Loop 1, results in a defect in Cr binding likely due to the impaired Loop 2 closure. In the presence of uci, stabilization of the loop in closed conformation may have enhanced Cr binding and catalytic activity in H61A MtCK. Similarly, uci may promote Cr binding and activity in other mutants with impaired Cr binding by enabling Loop 2 closure. Taken together, the combination of biochemical and structural data suggests uci may have a dual effect of competitive and uncompetitive mode of inhibition given its ability to bind MtCK regardless of presence or absence of substates, yet the functional effect of the binding is only seen through substrate Cr binding.

## Conclusion

In this work, we identified and characterized small-molecule inhibitors of mitochondrial creatine kinase, an enzyme whose increased activity is associated with diverse types of cancers. By integrating high-throughput screening, biochemical characterization, and cryo-EM structure determination, we uncovered mechanistically diverse MtCK modulators such as orthosteric and allosteric inhibitors. Structural elucidation of MtCK in complex with a newly discovered allosteric inhibitor with dual competitive and uncompetitive role revealed a previously unrecognized binding pocket and provided a molecular basis for its inhibitory mechanism, in which stabilization of a closed active-site loop restricted substrate access and product release. Together, these studies present chemical probes valuable for validating MtCK as a therapeutic target and establish a structural framework for future structure-guided optimization and rational design of next-generation MtCK inhibitors.

## Data and Code Availability

The cryo-EM maps of MtCK are deposited to the Electron Microscopy databank with accession numbers of EMDB-73720, -73766 and -73767. The model coordinates of MtCK structures are deposited in Protein Data Bank with PBD codes of 9Z0P, 9Z2D and 9Z2F. They are publicly available as of the date of publication.

## Acknowledgements

This work was supported by NIH/NCI grant (R01 CA251910) to Eduard Sergienko and Taro Hitosugi. Cryo-EM data was collected at Cryo-EM facility at Sanford Burnham Prebys Medical Discovery Institute supported by NIH/NCI grants (S10 OD026926 and P30 CA030199). MtCK1 protein production and MST were carried out at the Protein Production and Analysis Group supported by R01 CA251910 and NIH/NCI grant P30 CA030199. Merve Demir was supported by Conrad Prebys Foundation award number CPF #651 and R01 CA251910.

## Author Contributions

Conceptualization, M.D. and E.S. Investigation, M.D., L.K., C.M., P.G., N.P., S.P., S.A., Z.Z., L.F. and A.B. Writing-Original Draft, M.D. Writing-Review and Editing, M.D., J.Z. and E.S. Visualization, Y.L. and M.D. Funding Acquisition, T.H. and E.S. Supervision, J.Z., and E.S.

