## Supplementary material for "Uncompetitive Allosteric Inhibitor of Mitochondrial Creatine Kinase Prevents Binding and Release of Creatine by Stabilization of Loop Closure": Figure S1, Table S1

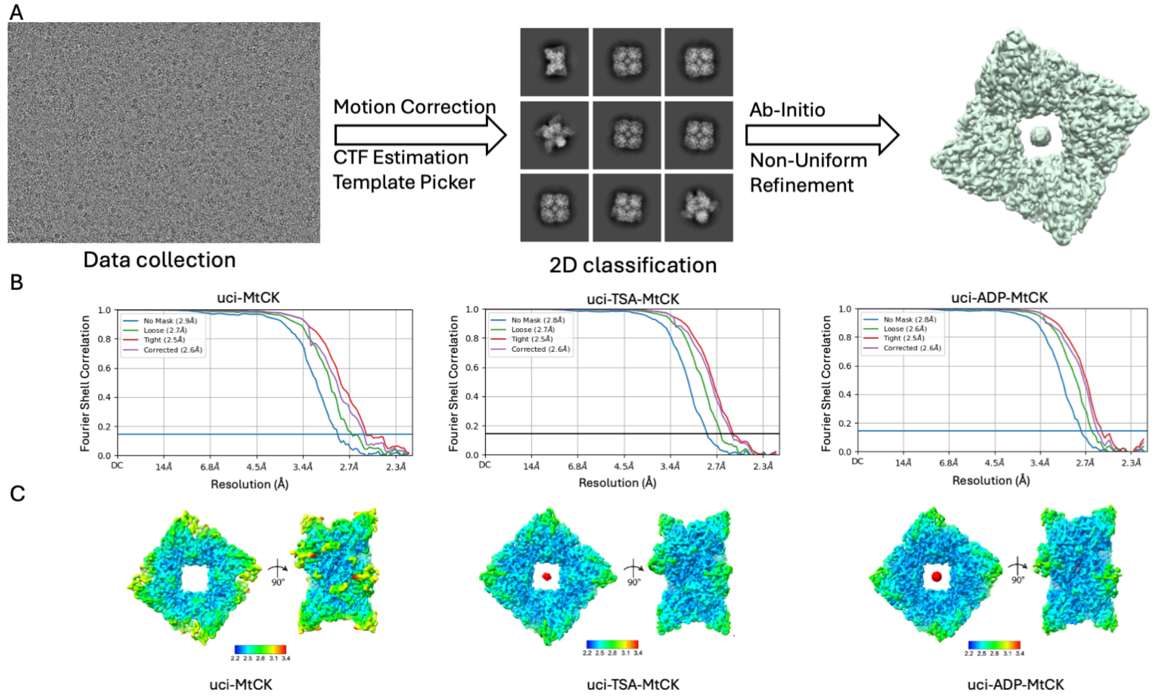


**Figure S1.** Cryosparc workflow for MtCK data processing. A) Collected movies were processed by motion correction, CTF estimation and particles picking using template pick. Particles were used for 2D classifications and to obtain the final maps. B) Fourier shell correlation (FSC) plots of each MtCK map. C) Final maps colored according to local resolution estimation by Cryosparc.

**Table S1. Cryo-EM data collection, refinement and validation statistics**.

| | uci-MtCK<br>EMDB-73720<br>PDB ID:9Z0P | uci-TSA-MtCK<br>EMDB-73766<br>PDB ID:9Z2D | uci-ADP-MtCK<br>EMDB-73767<br>PDB ID:9Z2F |
|---|---|---|---|
| **Data collection and processing** | | | |
| Voltage (kV) | 300 | 300 | 300 |
| Electron exposure (e/Å$^2$) | 34 | 34 | 34 |
| Defocus range (μm) | 0.9-2.1 | 0.9-2.1 | 0.9-2.1 |
| Pixel size (Å) | 1.064 | 1.064 | 1.064 |
| Symmetry imposed | D4 | D4 | D4 |
| Initial particle images (no.) | 4,227,185 | 3,529,024 | 2,731,259 |
| Final particle images (no.) | 581,677 | 1,083,440 | 1,213,097 |
| Map resolution (Å)<br>FSC threshold | 2.56<br>0.143 | 2.56<br>0.143 | 2.59<br>0.143 |
| **Refinement** | | | |
| Initial model used (PDB ID) | 9B05 | 9B14 | 9B14 |
| Model composition<br>Protein residues<br>Ligands | 2736<br>8 | 2944<br>24 | 2944<br>16 |
| R.m.s. deviations<br>Bond lengths (Å)<br>Bond angles (°) | 0.003<br>0.540 | 0.004<br>0.565 | 0.003<br>0.581 |
| Validation<br>MolProbity score<br>Clashscore<br>Poor rotamers (%) | 1.43<br>4.64<br>1.60 | 1.46<br>3.42<br>0.75 | 1.64<br>4.93<br>0.71 |
| Ramachandran<br>Favored (%)<br>Allowed (%)<br>Ouliers (%) | 97.83<br>2.17<br>0 | 95.29<br>4.47<br>0.24 | 94.36<br>5.40<br>0.24 |
